# Embryonic exposures to flame retardant tetrabromobisphenol A (TBBPA) disrupts dorsoventral patterning in zebrafish

**DOI:** 10.1101/2024.12.16.628760

**Authors:** Kanchaka Senarath Pathirajage, Rosemaria Serradimigni, Sunil Sharma, Subham Dasgupta

## Abstract

Tetrabromobisphenol A (TBBPA), a widely used flame retardant in commercial products such as synthetic textiles, plastics, and electronics poses potential toxicity risks through indoor exposure. This study aims to leverage zebrafish as a model to study TBBPA impacts on zebrafish dorsoventral patterning—a process that lays the foundation of an embryo’s axial determination and localization of specific tissues and organs. Zebrafish embryos were exposed to varying concentrations of TBBPA (0-10 µM) at either 0.75- or 6-hours post-fertilization (hpf) and phenotyped at 8 or 24 hpf. Following this, whole-mount immunohistochemistry (IHC) was conducted out to quantify various proteins important in the BMP signaling pathway, epithelial-to-mesenchymal transition (EMT) and ectoderm and mesoderm germ layers. Importantly, these assessments were done at environmentally relevant concentrations ranging down to nM and pM levels. Our data showed a significant concentration-dependent increase in ventralization phenotypes, marked by enlarged blood island area, coupled with a disruption of the ventral-to-dorsal gradient of pSMAD protein levels, indicating BMP signaling disruptions. Perturbations in epithelial-to-mesenchymal transition (EMT), as evidenced by changes in E- and N-cadherin expression and Snail2 transcription factor levels, indicated impaired cell migration on TBBPA exposures. We then interrogated if TBBPA impacts germ layers and structures derived from specific germ layers. We observed significant concentration-dependent changes in levels of Sox2 and Sox10- both indicators of neural crest cell formation and differentiation from the ectoderm. We also observed a significant reduction of Tbx16- a marker of paraxial mesodermal cells. Collectively, both these data show TBBPA-induced impact on germ layers. Finally, we examined specific cell types derived from ectoderm and mesoderm and showed TBBPA-induced inhibition of cartilage development (derived from ectodermal neural crest cells) and blood cell development (derived from mesodermal cells). Our findings collectively demonstrate that TBBPA-induced disruptions in early developmental signaling and dorsoventral patterning may contribute to systemic toxicity in zebrafish embryos. Importantly, we see these disruptions at environmentally relevant concentrations, reinforcing the importance of continued interrogation of TBBPA in targeting early embryogenesis.

## Introduction

Tetrabromobisphenol A (TBBPA) is a brominated flame retardant that has been extensively used in commercial products like textiles, car seats, polystyrene foams, furniture, epoxy resins and electronic devices to reduce the flammability (Miao et al., 2023). The overall production quantity of TBBPA in 2015 ranged from 50 million to 100 million pounds (EPA.,2020). Because of its high production volume, the risk of having exposure to TBBPA through ingestion, respiratory system, or dermal absorption is significant (Miao et al., 2023). Prior studies have shown evidence of TBBPA-induced endocrine disorders, tumor incidences, developmental defects, neurotoxicity, immunotoxicity, and reproductive toxicity even at environmental-related lower doses (S. Li et al., 2023).

This study focuses on TBBPA-induced impacts on dorsoventral patterning-a crucial developmental event that determines the body axis. Developmental effects of TBBPA are a concern because of their substantial detection in cord blood (Cariou et al., 2008), breast milk (Shi et al., 2009), placental tissue (Alzualde et al., 2018) as well as maternal milk and cord blood at nM levels (Zhao et al., 2022). In fact up to 3.65 µg/L (6.710 μM) of TBBPA has been detected in human plasma which was collected from 140 voluntary donors who are registered doctors and nurses (Ho et al., 2017). Dorsoventral patterning is a sensitive window of development, constituting a coordinated series of cell migrations and differentiations to organize cells along the body axis preceding organogenesis(Pang et al., 2022). This process is regulated by the bone morphogenetic protein (BMP) signaling; a gradient of BMP signaling in the blastomere as well as in germ layers that determines the fate of the cells (Pomreinke et al., 2017). This BMP gradient delineates the development of structures, such as the head from dorsal regions with low BMP signaling, form posterior structures, like the tail or coccyx, in ventral regions with high BMP signaling (Pomreinke et al., 2017). Additionally, mobility of cells and their migration along the body axis is regulated by altered levels of cell adhesion proteins, constituting epithelial to mesenchymal transition (EMT). Drugs or environmental chemicals that interfere with any of these signaling mechanisms or developmental events can trigger adverse impacts on embryogenesis. For example, disruption of BMP signaling has been contributed to the decreased intrinsic repair capacity of damaged cartilage (Deng et al., 2018). Likewise, a disruption of EMT has been implicated in the ectoderm, an increase in epidermal cells indicated a fate shift from neural crest to epidermal identity, while some neuroectodermal cells changed to neural crest cells, resulting in a notable decline in brain cell numbers (Pang et al., 2022). Therefore, studying impacts of xenobiotics on these processes is much needed to examine the molecular factors they target and interrogate mechanisms of developmental toxicity.

Zebrafish embryos has been used as the animal model for this project because of the high reproductive capacity, rapid and external development, transparency of embryo, easy to manipulate embryonically, used for chemical screens and high (>75%) similarity with the human genome thus making zebrafish an excellent model for vertebrate developmental biology (Veldman and Lin, 2008).Phenotyping of developmental defects in zebrafish has been widely used to identify early signs of toxicity that focus on developmental signaling pathways (Dasgupta et al., 2021). For example, zebrafish embryos treated with another flame retardant, tris(1,3-dichloro-2-propyl) phosphate (TDCIPP), showed gastrulation and dorsoventral patterning defects that was linked to disruption of mesodermal cell localization and somitogenesis (Dasgupta et al., 2018). Another study that investigated early developmental defects in zebrafish embryos exposed to the ubiquitous pollutant bisphenol A (BPA), three stages (60–75% epiboly, 8–10 somite, and prim-5), revealing that BPA exposure significantly altered early dorsoventral patterning, segmentation, and brain development within 24 hours (Liu et al., 2021). Within our study, we leverage similar phenotypic strategies to interrogate how TBBPA targets dorsoventral patterning. We first identify TBBPA-induced phenotypes and conduct a series of whole mount immunohistochemistry experiments at environmentally relevant concentrations to examine levels of proteins associated with BMP signaling, germ layers and EMT. We then conduct follow-up studies of cartilage and blood development to examine how disruptions of dorsoventral patterning can affect formation of downstream structures.

## Methodology

### Zebrafish rearing and embryo collection

Specific pathogen-free 5D wild type zebrafish were reared at a maximum density in non-chlorinated water glass tanks at a temperature of 27 ± 1 ° C and a photoperiod of 14:10 h (light/dark) and were fed with artificial feed three times a day in Aquatic Animal Research Laboratory at Clemson University. *sox10*:eGFP fish were received as a gift from Dr. Rosa Uribe at Rice University and maintained in a similar manner. Embryos were transferred to petri dishes (∼30–50 embryos in a 100-mm Petri dish) containing embryo medium (5 mM sodium chloride, 0.33 mM calcium chloride, 0.33 mM magnesium sulphate and 0.17 mM potassium chloride) by diluting 1 in 60 with distilled water). All adult zebrafish breeders were handled per an Institutional Animal Care and Use Committee-approved animal use protocol (No. 20220434) at Clemson University.

### Chemicals

TBBPA (97 % purity) (CAS #: 79–94-7) was purchased from Sigma-Aldrich. A stock solution was prepared using high-performance liquid chromatography-grade dimethyl sulfoxide (DMSO) and stored in 20 mL glass vials sealed with parafilm at room temperature. Prior to each experiment, working solutions were prepared using embryo media by first establishing a stock working concentration and then performing serial dilutions to obtain the desired concentrations.

### Chemical exposure and phenotyping

Embryos were exposed either at 0.75 h post-fertilization (hpf) or 6 hpf to 2 mL of TBBPA solution (0.1% DMSO as vehicle) of 0, 0.625, 1.25, 2.5, 5, 10, 20 µM with 3 replicates and 10 embryos per replicate in 10 mL glass beakers. Exposures were conducted in complete darkness. At 24 hpf, embryos were observed for ventralization phenotypes and grouped based on Dasgupta et al 2021.

### Whole-mount immunohistochemistry (IHC)

Embryos were exposed to TBBPA (0, 0.00005, 0.05, 0.5 or 5 µM) at 0.75 or 6 hpf and collected for IHC at 8 or 24 hpf. Briefly, embryos were fixed overnight in 4% paraformaldehyde (PFA), dechorionated and immunohistochemistry was conducted according to prior protocols (Dasgupta et al., 2021) with the following antibodies: anti-goat pSMAD 1/5/9 (1:100, Cell Signaling), anti-E-cadherin (RR1; 1:2.5; Developmental Studies Hybridoma Bank/ DSHB), N-cadherin (6B3;1:2.5, DSHB), anti-Snail2 (IE6; 1:2.5; DSHB), anti-Sox2 (1:100,Abcam,anti-VegT (1:10; ZIRC), anti-Sox10 (1:200, Abcam). After washing, embryos were counter stained in corresponding Alexa Fluor antibodies (1:500; Sigma) and visualized on an Echo Revolve Microscope. Images were analyzed and quantified on Image J and statistical estimations were conducted in GraphPad Prism 10.

### Alcian Blue cartilage staining

Embryos were exposed to 0, 0.0005, 0.5 or 1 µM TBBPA from 0.75 hpf to 6 days post fertilization (dpf) and fixed in 4% PFA for 2 hours at 4 °C, followed by washing with PBS. Embryos were then dehydrated in 50 % ethanol for 10 minutes. Following this, cartilage was stained with 0.4% Alcian Blue 8GX (Sigma-Aldrich, St Louis, Missouri) in 80 mM MgCl_2_ and 50% ethanol as described in (Dasgupta et al., 2023) and kept overnight in a shaker at room temperature. Pigmentation was removed by incubating fixed larval samples for 30 min in a mixture of 3% H_2_O_2_ and 1% KOH. Following this, embryos were incubated in clearing solutions containing 25% glycerol followed by 50% glycerol in successive overnight incubations, both in 0.25% KOH solution. Larvae were imaged using an Olympus TH4–100 equipped with a Lumenera Infinity 8–8 camera (Ottawa, Ontario, Canada) at 4X magnification. Images were analyzed and quantified on Image J and statistical estimations were conducted in GraphPad Prism 9. for lower jaw length (LJL), intercranial distance (ICD), ceratohyal cartilage length (CCL) and area of the cartilage.

### O-dianisidine staining for hemoglobin

Embryos were exposed to 0, 0.00005, 0.005 or 0.5 µM TBBPA from 0.75 hpf to 3 (dpf), and hatched embryos were stained in the dark for 30 minutes at room temperature within a solution containing o-dianisidine (0.6 mg/mL, 0.01 M sodium acetate (pH 4.5), 0.65% H_2_O_2_, and 40% (vol/vol) ethanol. Once stained, embryos were transferred into 1.5mL Eppendorf tubes and washed with molecular grade water 3 times, and then fixed in 1mL of 4% paraformaldehyde for at least 1 h at room temperature. Pigments were removed from fixed embryos by incubating with a depigmentation solution of 0.8% KOH, 0.9% H_2_O_2_, and 0.1% Tween-20 for 30 minutes at room temperature and were shaken every 10 minutes. Embryos were then washed with phosphate-buffered saline (PBS) and then fixed again in 4% PFA overnight in PBS at 4°C. All embryos were oriented in dorsal recumbency and imaged using an Olympus TH4–100 equipped with a Lumenera Infinity 8–8 camera (Ottawa, Ontario, Canada) at 4X magnification. Images were analyzed using a color range tool within Adobe Photoshop. Sample colors were selected within vehicle control images to represent the o-dianisidine stain, and then control color samples were used to measure the stained area. Images were analyzed and stained area was quantified on Adobe Photoshop.

### Statistics

All statistical estimations were done in Graphpad Prism 9. Statistical significance was determined using a one-way ANOVA, followed by Dunnett’s post hoc tests for comparison of each concentration to the DMSO controls (p<0.05). For ventralization phenotyping, number of normal embryos in each replicate was used to estimate differences. Detailed results from statistical tests (ANOVA p values and Dunnett’s p values) are included within each figure.

## Results and Discussion

Dorsoventral patterning lays the foundation for axial determination of an embryo and localization of specific tissues and organs (Langdon and Mullins, 2011). Despite prior studies investigating the TBBPA-induced developmental toxicity of zebrafish embryos (Chen et al., 2016) (Zhu et al., 2018), we are the first to assess these dorsoventral patterning markers at environmentally relevant concentrations (nM or pM levels). Notably, we examine EMT-a process that is not typically explored in developmental toxicology at an *in vivo* level. For this study, we pursued several different exposure scenarios to examine various biological windows, wherever possible. By initiating exposures at either 0.75 or 6 hpf, we sought to study and compare if pre-pluripotency signals, prior to 6 hpf, play a differential role in TBBPA-induced effects on dorsoventral patterning. Likewise, by sampling embryos at 8 or 24 hpf, we sought to examine impacts both at the onset of dorsoventral patterning/ EMT (8 hpf) and during these processes when the anteroposterior polarity has already been established and some rudimentary organs have already been formed (24 hpf). Our work bridges the gap in understanding the molecular targets of TBBPA that potentially drive dorsoventral patterning defects, focusing on the bone morphogenetic protein (BMP) signaling and EMT.

### TBBPA causes ventralization

Embryonic exposure to a range of TBBPA concentrations show a concentration-dependent increase in ventralization (ANOVA p<0.0001; **Figure 1A and Figure 1B)**, noted by expansion of ventral areas such as blood island (ANOVA p<0.0001; **Figure 1C)**. The different intensities of ventralization (V1 to V4) indicate the strength of effects on dorsoventral patterning. Notably, we see this effect regardless of when we initiate the exposures-0.75 **(Figure 1A)** or 6 hpf **(Figure 1B)**. Interestingly, impacts of the 6 hpf initiation experiments are stronger, with stronger ventralization intensities at higher concentrations **(Figure 1B)**-the roots of this difference in effects need to be further interrogated. Consistent with this, a prior study showed that exposure to TBBPA resulted in an increased number of embryonic cells adopting a ventral cell fate (Pang et al., 2022). These phenotypes led us to hypothesize the interference of the bone morphogenetic protein (BMP) signaling pathway and EMT (Wang et al., 2014), both of which regulate dorsoventral patterning. Disrupting the BMP signaling pathway and epithelial-to-mesenchymal transition can significantly affect zebrafish development, resulting in abnormal body axis formation and impairing the development of vital organs like the heart and kidneys (Mahadeva et al., 2025). This can also interfere with somite formation, impacting muscle and skeletal growth (Kim et al., 2014; Nifuji et al., 1997). Additionally, altered cell migration may prevent proper tissue organization, potentially increasing embryonic mortality (Gomez-Puerto et al., 2019); surviving embryos might experience behavioral issues or physical deformities as adults, affecting their overall health and reproductive success (Gomez-Puerto et al., 2019).

**Figure 1.**
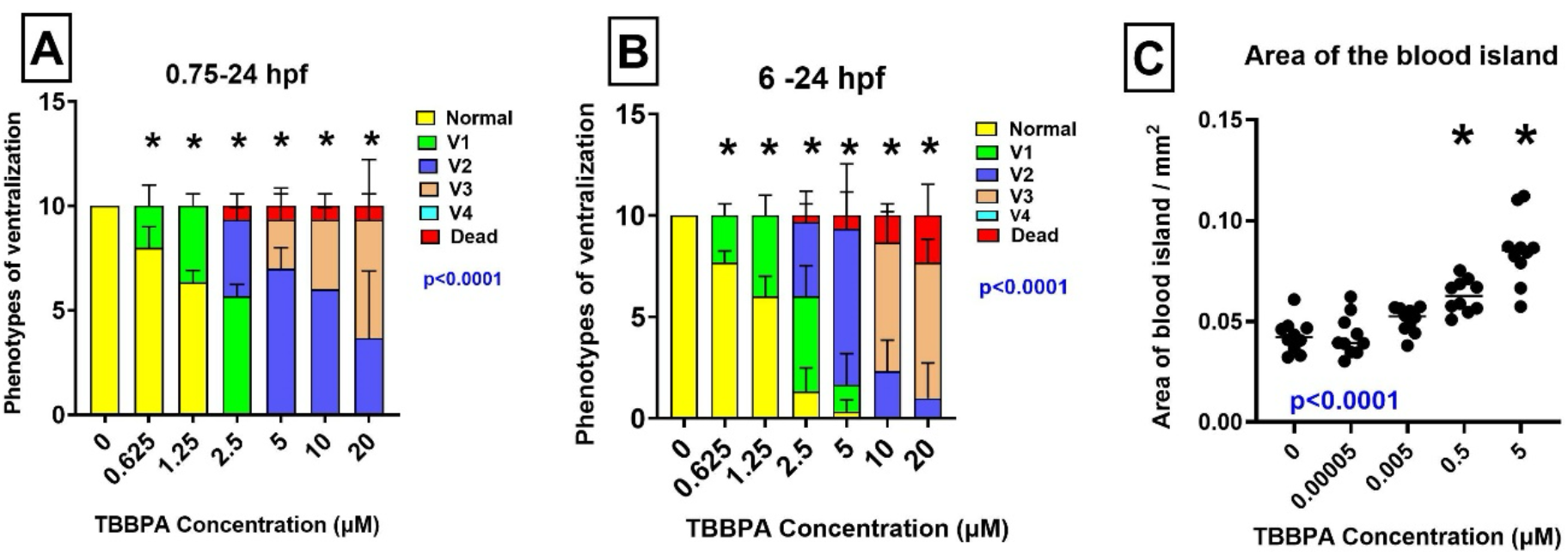
TBBPA ventralizes zebrafish embryos. (A) 0.75-24 hpf exposure. (B) 6-24 hpf exposure. (C) Area of the blood island. One-way ANOVA p values are denoted by blue text; asterisk (*) denotes statistically different (p<0.05) from 0 µM based on Dunnett’s post hoc test following 1-way ANOVA

### TBBPA disrupts BMP signaling

Phospho-smad (pSMAD) proteins are suitable indicators of BMP signaling and show a ventral-to-dorsal gradient during gastrulation (Ramel and Hill, 2013). We conducted IHC for pSMAD following two exposure scenarios-0.75-8 and 6-8 hpf. Our data showed two related outcomes. First, there were significant changes in pSMAD gradient across the exposures **(Figure 2A and Figure 2B)**, with statistically significant differences down to 500 nM. And second there was an overall increase in BMP signaling across the embryo, denoted by increased pSMAD fluorescence **(Figure 2C and Figure 2D)**. Taken together, these indicate increased BMP signaling and loss of BMP gradient, that can have significant repercussions for dorsoventral patterning. This data is also consistent with altered mRNA levels of BMP signaling inhibitors *chordin* and *sizzled*, as seen in our previous data on mRNA seq on 0.75 - 5 hpf of TBBPA exposure on zebrafish **(Table 1)** (Serradimigni et al., 2024). Importantly we see this inhibitory effect only after 2 hr of exposure (6-8 hpf) during gastrulation. Taken together, these imply that TBBPA targets specific BMP-associated targets during gastrulation that impacts the BMP signaling gradient.

**Figure 2.**
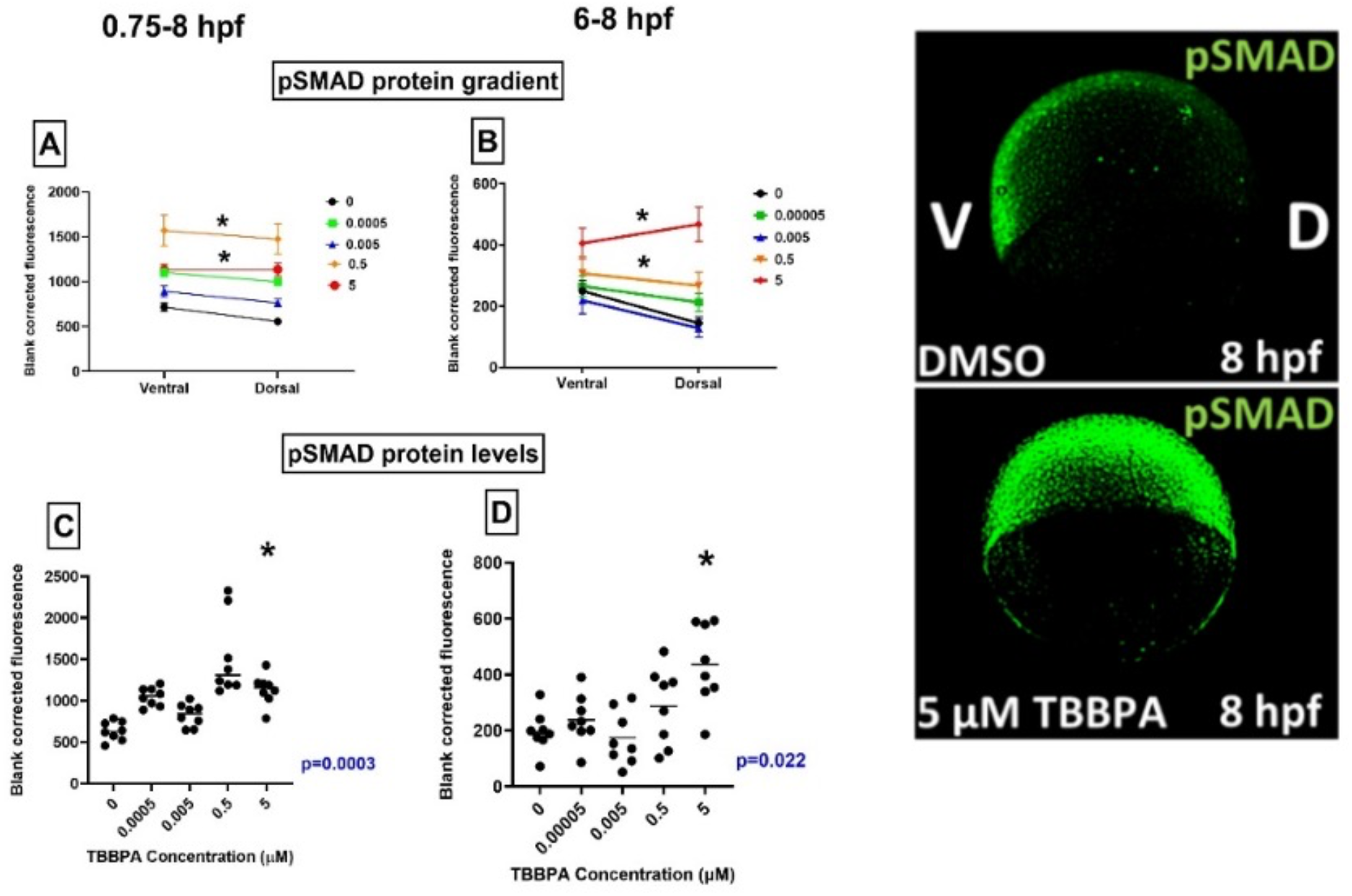
TBBPA disrupts pSMAD gradient. (A) 0.75-8 hpf exposure. (B) 6-8 hpf exposure. representative images on the right. Overall, 1-way ANOVA p values are denoted by blue text; asterisk (*) denotes statistically different (p<0.05) from 0 µM based on Dunnett’s post hoc test following 1-way ANOVA.

**Table 1.** RNA seq data of BMP signaling genes on 0.75 - 5 hpf TBBPA based exposures based on Serradimigni et al 2024

| Gene | log2FC |
| --- | --- |
| *ved* | -2.71 |
| *vox* | -2.18 |
| *bambia* | -2.12 |
| *bmp2b* | -1.2 |
| *bmp2a* | -2.8 |
| *bmp4* | -3.3 |
| *bmp7a* | -2.9 |
| *smad7* | -2.4 |
| *szl* | -3.58 |
| *chrd* | -3.9 |

### TBBPA disrupts factors driving epithelial-to-mesenchymal transition (EMT)

Epithelial-to-mesenchymal transition (EMT) is a critical process for cell migration and establishment of cell types and body structures that causes epithelial cells to lose their epithelial characteristics and acquire mesenchymal characteristics (Kalluri and Weinberg, 2009). This process is largely regulated by BMP signaling as well as several EMT transcription factors that, in turn, modulate cell adhesion proteins and other biomolecules (Wang et al., 2014). To study this process, we conducted IHC on epithelial marker cell adhesion protein E-cadherin, mesenchymal marker N-cadherin and EMT transcription factor Snail2. Our prior sequencing data showed that TBBPA reduced transcript levels of these factors at 5 hpf **(Table 1)**. During EMT, the EMT transcription factors facilitate the inhibition of E-cadherin and increase of N-cadherin within cells, triggering the migratory process. We initiated exposures at 0.75 or 6 hpf and sampled embryos at 8 or 24 hpf. While exposure initiation did not make a difference in the trend of response, responses somewhat varied based on the developmental stage we examined them at.

#### 8 hpf

E-Cadherin levels showed an overall concentration-dependent increase for both 0.75-8 and 6-8 hpf treatments (p=0.02 and <0.001 respectively), with significant increases in environmentally relevant concentrations of 0.005 µM **(Figure 3A and Figure 3B)**. Interestingly, shorter exposure (6-8 hpf) led to more extensive impacts compared to longer exposures-an outcome that may be attributed to metabolism of TBBPA and needs to be examined further. Along with increased E-cadherin at 8 hpf, N-cadherin **(Figure 3C and Figure 3D)** and Snail2 levels **(Figure 3E and Figure 3F)** were reduced (p<0.0001 in each case). Collectively, these suggest that at the onset of EMT, TBBPA inhibits the EMT process by inducing anti-migratory E-cadherin and inhibiting pro-migratory N-cadherin and these may be driven by targeted inhibition of EMT transcription factors such as Snail2. Importantly, these effects are seen at environmentally relevant, nM level concentrations. This inhibition of cell migration may be causative behind downstream dorsoventral patterning defects, which is reliant on coordinated cell migration along the body axis (Pang et al., 2022).

**Figure 3.**
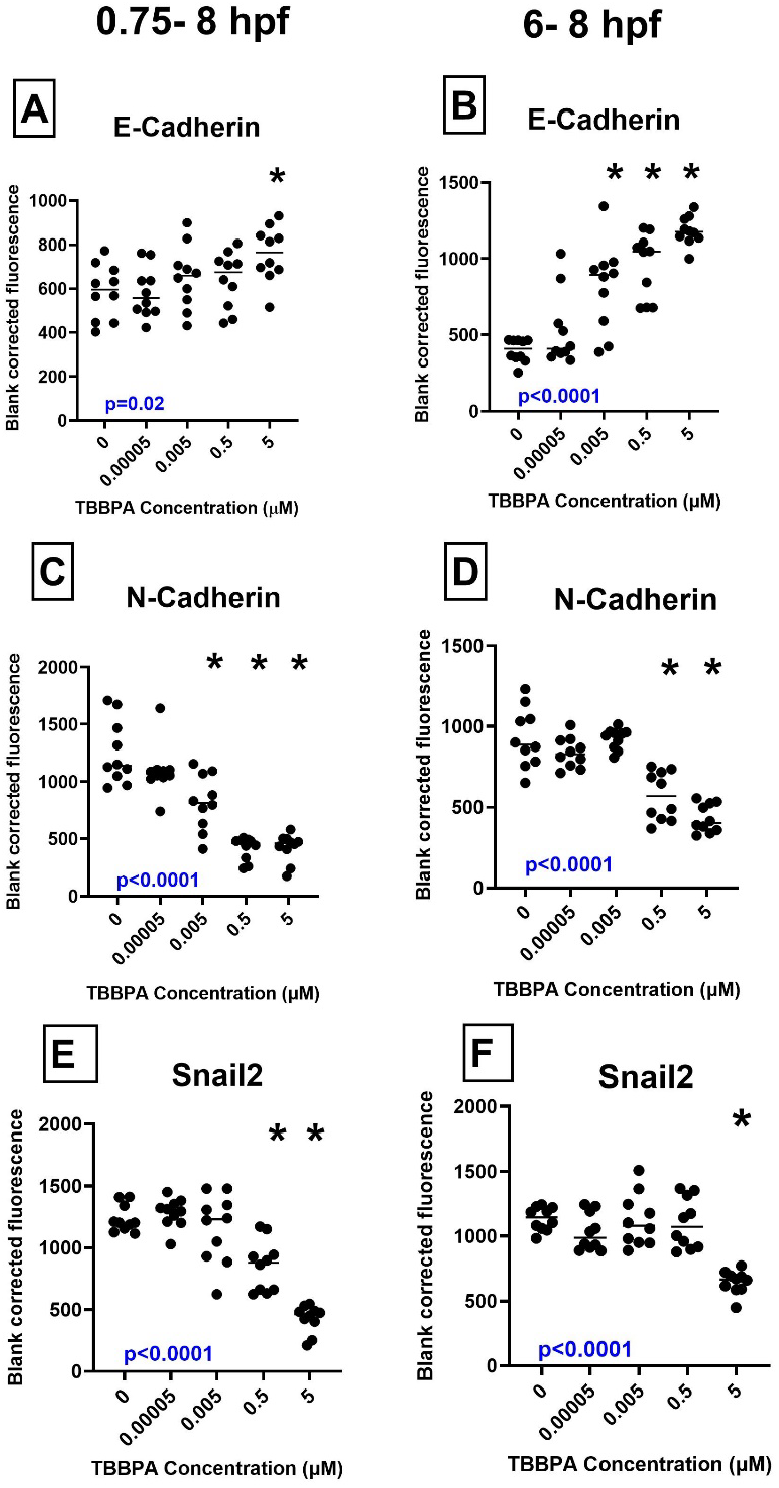
TBBPA alters EMT factors at 8 hpf **(A, B)** E-Cadherin **(C, D)** N-Cadherin **(E, F)** Snail2. Exposure durations were from 0.75-8 or 6-8 hpf as indicated above the figures. Overall, 1-way ANOVA p values are denoted by blue text; asterisk (*) denotes statistically different (p<0.05) from 0 µM based on Dunnett’s post hoc test following 1-way ANOVA

#### 24 hpf

Examination of EMT factors at 24 hpf presented a somewhat different scenario. For E-cadherin, expectedly, 0.75-24 hpf experiments showed overall increases (p<0.0001) **(Figure 4A)**, but 6-24 hpf showed increases at only the top concentration **(Figure 4B)**. The lower levels of E-cadherin in zebrafish exposed to TBBPA from 0.75 to 24 hpf, compared to 0.75 to 8 hpf may be due to the prolonged exposure from 0.75-24 hpf that may cause cumulative cellular stress. This, in turn, may disrupt key signaling pathways like Wnt or TGF-β pathways that promote E-cadherin expression leading to further downregulation of E-cadherin (Ogata et al., 2007). As expectably N-cadherin levels were reduced both 0.75-24 hpf (p<0.0001) **(Figure 4C)** and 6-24 hpf (p<0.001) **(Figure 4D)** exposures. Snail2 levels were decreased in 0.75-24 hpf exposure (p<0.0001) **(Figure 4E)**, but the concentration response resembled a U-shaped curve for 6-24 hpf **(Figure 4F)**. Such response may be associated with compensatory genetic responses at higher TBBPA concentrations.

**Figure 4.**
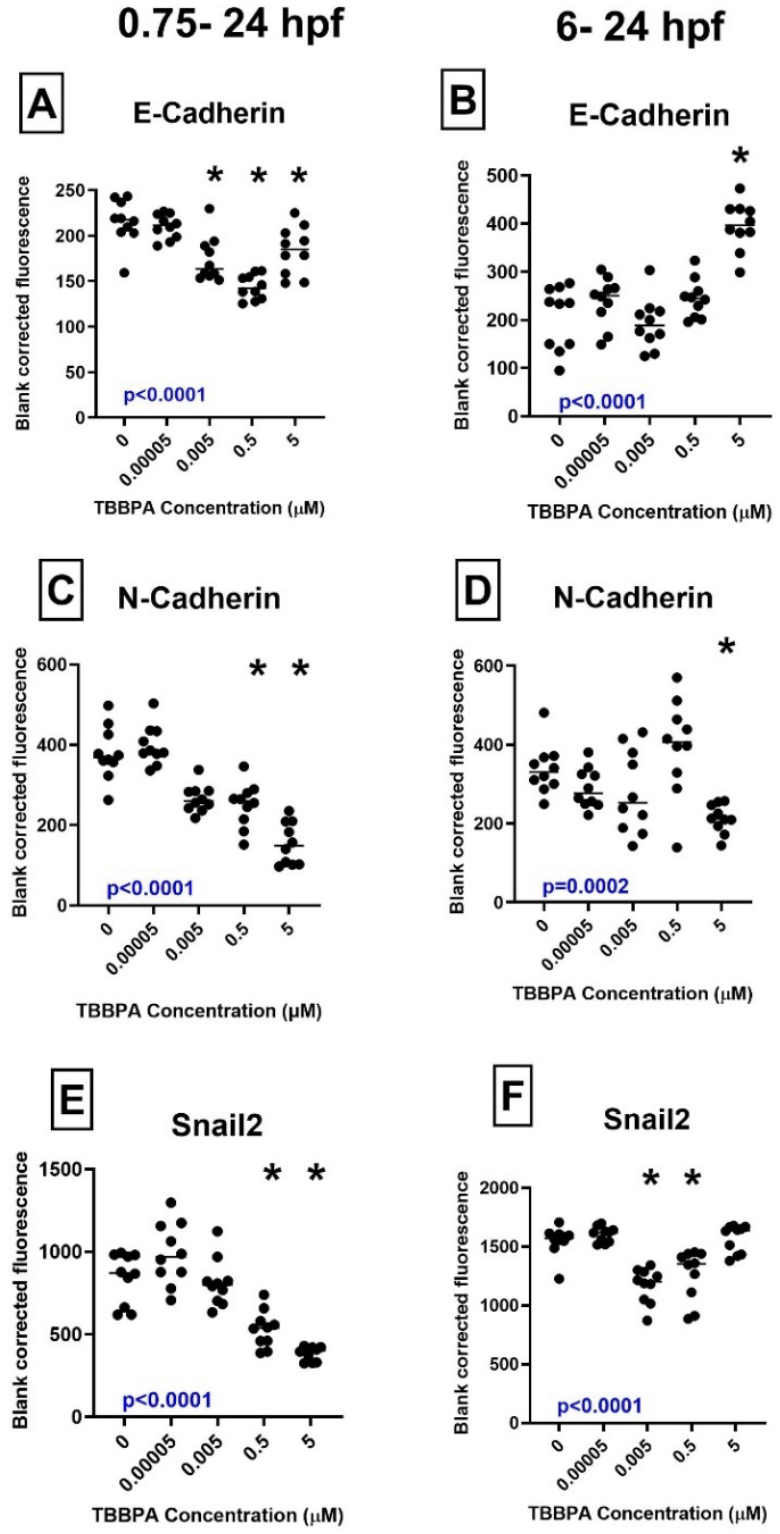
TBBPA alters EMT factors at 24 hpf. (A, B) E-Cadherin (C, D) N-Cadherin (E, F) Snail2. Exposure durations were from 0.75-24 or 6-24 hpf as indicated above the figures. Overall, 1-way ANOVA p values are denoted by blue text; asterisk (*) denotes statistically different (p<0.05) from 0 µM based on Dunnett’s post hoc test following 1-way ANOVA.

Overall, these data show that TBBPA targets EMT factors and potentially inhibits this process. Importantly, the EMT process has been shown to be dependent on BMP signaling (Wang et al., 2014), providing a basis for studying BMP-EMT interactions for TBBPA treatments. Consistent with this, another study found out that in human colorectal cancer, Snail1 depends on BMP signaling with SMAD4 transcription factor for EMT execution, identifying a BMP-dependent gene signature that regulates EMT (Frey et al., 2020). Additionally, EMT is a critical process for cancer metastasis and multiple compounds with anticancer activity have been shown to inhibit EMT by suppressing key molecules or pathways such as E and N-cadherin (Loh et al., 2019), providing a potential link between carcinogenesis and TBBPA.

### Germ layer-specific effects-ectoderm-derived neural crest cells and head cartilage

We further investigated TBBPA-induced impacts on the two germ layers-ectoderm and mesoderm, as well as select tissues derived from these germ layers. Ectoderm is a germ layer that differentiates into parts of the central nervous system, cartilage (neural ectoderm), and other structures such as hair and skin (Koehler et al., 2014). The differentiation of the ectoderm into its target tissues is largely moderated by BMP signaling gradient; high BMP regions give rise to skin and notochord, intermediate level of BMP regions give rise to neural crest cells and low BMP regions give rise to neural tube and then the brain (Koehler et al., 2014). We examined TBBPA-induced impacts on-1) neural crest cells and neural crest cell-derived cartilages and 2) the developing brain. First, we used Sox10-a marker of late-stage neural crest cells-to interrogate the presence of neural crest cells. To study the levels of Sox10, we conducted both IHC using an anti-Sox10 antibody and a *sox10*:eGFP reporter fish line. Both data show a decrease in levels of Sox10 within the developing head, indicating a reduction of cranial neural crest cells at 0.75 hpf (p = 0.0026) **(Figure 5A)** and 6 hpf (p = 0.0452) **(Figure 5B)**. In addition, we also saw a decrease in Sox2 (p<0.001)-a marker of pluripotency **(Figure 6A-D)**; notably, Sox2 is also known as a rheostat for neural crest cells, maintaining its flow and number in the head(Klymkowsky et al., 2010). Collectively, these indicate that TBBPA impacts pluripotency and neural ectoderm-a fact that is consistent with our previous work that shows TBBPA-induced effects on maternal-to-zygotic transition (Serradimigni et al., 2024). Notably, similar to EMT markers, we see these effects on these molecular markers at sub-µM level environmentally relevant concentrations. We also morphometrically quantified the midbrain-hindbrain barrier at 24 hpf and found reductions in height of mid brain and hindbrain barrier (p<0.0001) **(Figure 7C)**, which indicate disruptive impacts on neural tube/brain.

**Figure 5.**
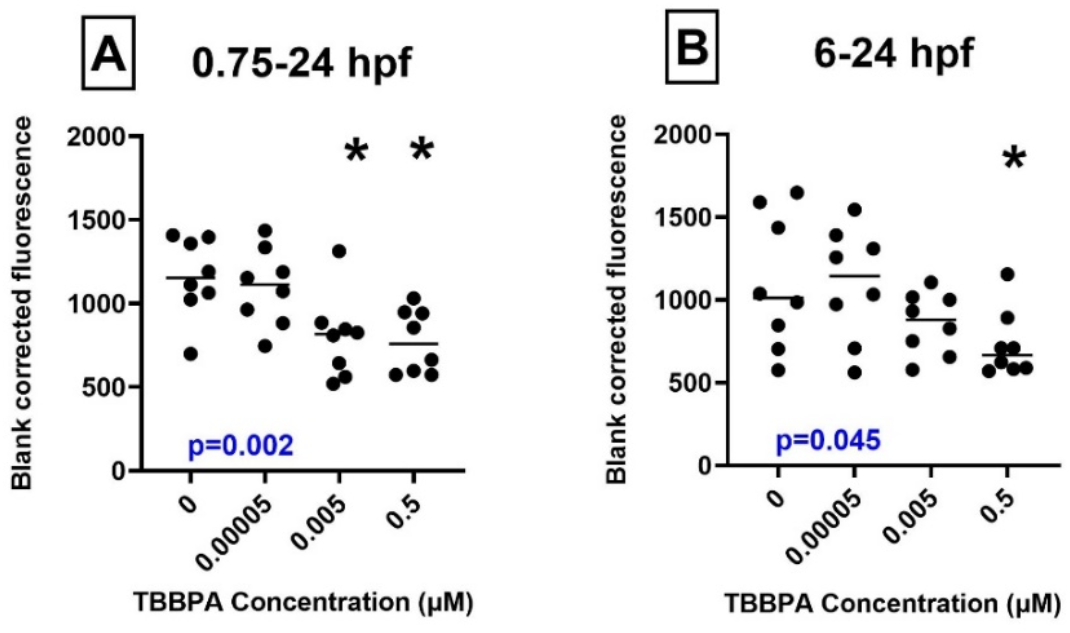
TBBPA reduces Sox10 levels. (A) 0.75-24 hpf and (B) 6-24 hpf. Overall, 1-way ANOVA p values are denoted by blue text; asterisk (*) denotes statistically different (p<0.05) from 0 µM based on Dunnett’s post hoc test following 1-way ANOVA.

**Figure 6.**
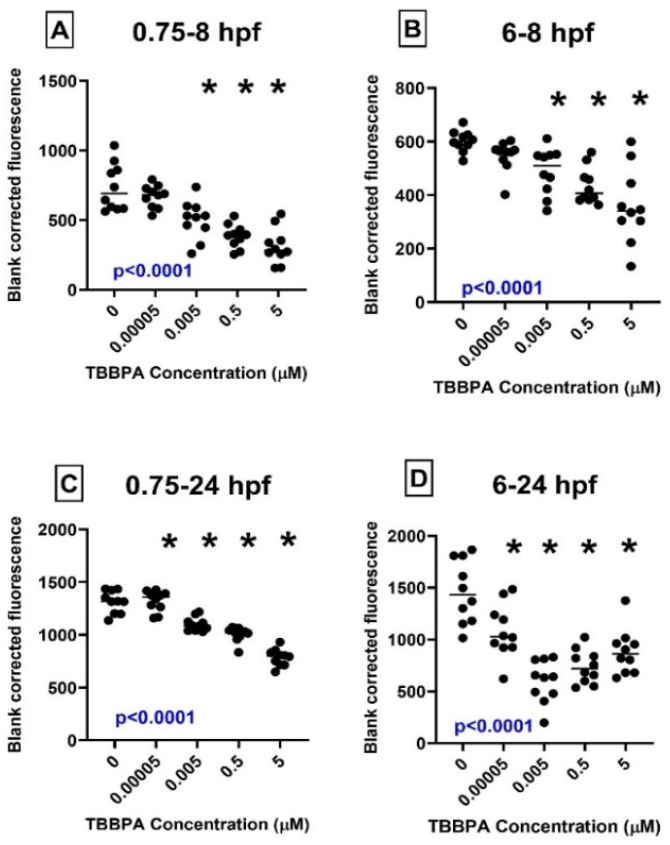
TBBPA alters Sox2 levels. (A) 0.75-8 hpf exposure. (B) 6-8 hpf exposure. (C) 0.75-24 hpf exposure. (D) 6-24 hpf exposure. Overall, 1-way ANOVA p values are denoted by blue text; asterisk (*) denotes statistically different (p<0.05) from 0 µM based on Dunnett’s post hoc test following 1-way ANOVA.

**Figure 7.**
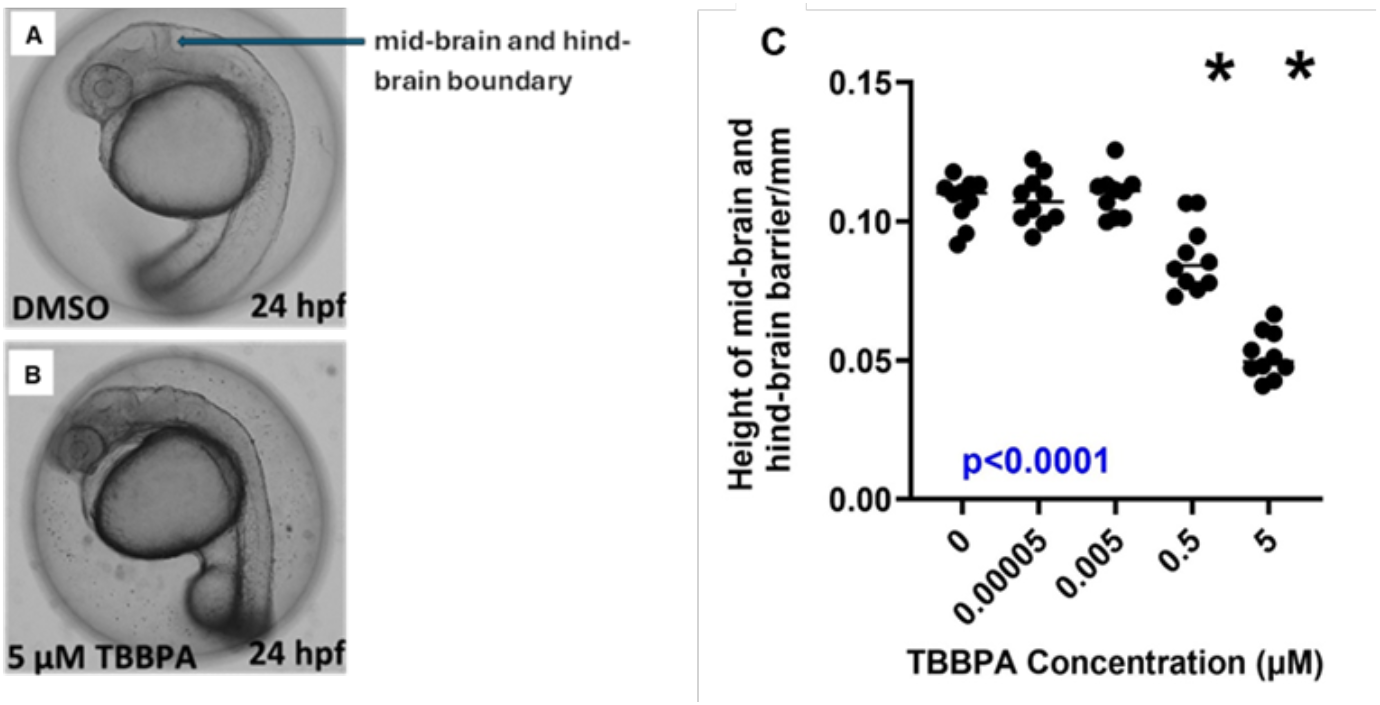
TBBPA decreases the height of the mid brain and hindbrain barrier. (A) Embryo exposed to DMSO. (B) Embryo exposed to 5 µM TBBPA (C) Heights of mid-brain and hind-brain barrier of embryos exposed to different TBBPA concentrations. Overall, 1-way ANOVA p values are denoted by blue text; asterisk (*) denotes statistically different (p<0.05) from 0 µM based on Dunnett’s post hoc test following 1-way ANOVA.

Cartilage is primarily derived from neural crest cells, which migrates from the neural tube to contribute to the formation of bone and cartilage in the craniofacial region (S. Li et al., 2023). The proper differentiation and migration of these neural crest cells are essential for the development of cartilaginous structures, such as the lower jaw, ceratohyal cartilage, and intercranial cartilage (Z. Li et al., 2023). As reductions in the pluripotency of embryonic stem cells and a decrease in the number of neural crest cells were observed, we hypothesized that exposure to TBBPA may disrupt the normal processes underlying cartilage formation. Therefore, we conducted Alcian Blue staining to measure head cartilage structures at 6 dpf. Our data showed that lower jaw length (p<0.0001) **(Figure 8A)**, ceratohyal cartilage length (p=0.009) **(Figure 8B)**, intercranial distance (0.0136) **(Figure 8C)** and the area of the cartilage (p<0.0001) (**Figure 8D)** were decreased significantly with TBBPA concentration specially both in 1 µM and in 0.5 µM. These findings indicate that improper differentiation of neural crest cells from the neural tube may play a significant role in causing abnormalities in brain development and may disrupt the neural crest cell-derived developing cartilage structures at 6 dpf.

**Figure 8.**
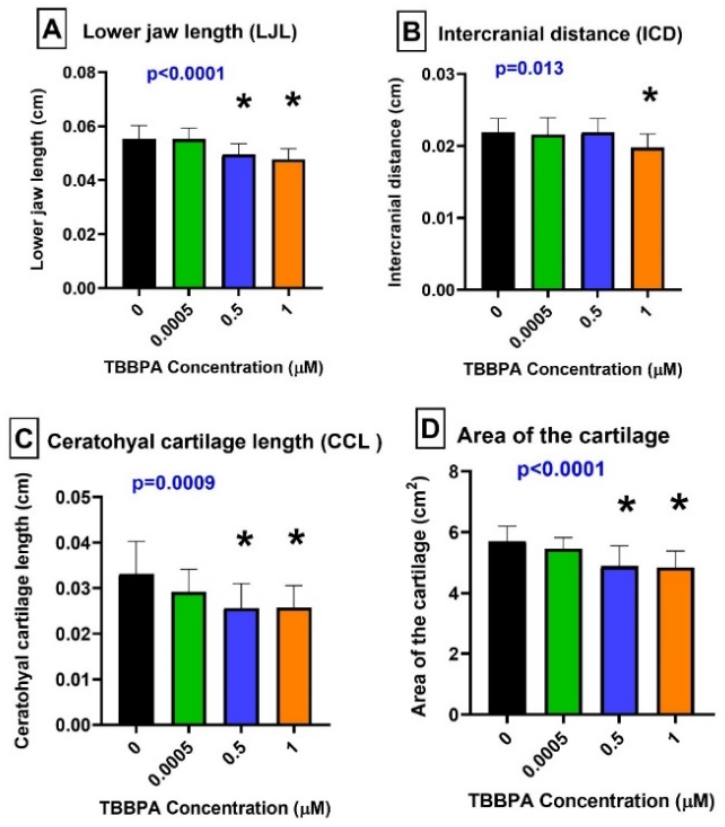
TBBPA alters cartilage development at 0.75 hpf – 6 dpf. (A) Lower jaw length (LJL). **(B)** Intercranial distance (ICD). (C) ceratohyal cartilage length (CCL**).(D)** Area of the cartilage. Overall, 1-way ANOVA p values are denoted by blue text; asterisk (*) denotes statistically different (p<0.05) from 0 µM based on Dunnett’s post hoc test following 1-way ANOVA

### Germ layer-specific effects-mesoderm and mesoderm-derived blood cells

We also sought to interrogate if TBBPA impacts mesoderm-another germ later that differentiates into muscles and blood cells. The *tbx16* gene (also known as T-box transcription factor 16) is involved in the formation of mesodermal tissues that give rise to various structures including muscles, the circulatory system, and bones located around the notochord during embryonic development (Morrow et al., 2017). We leveraged Tbx16 as a marker of mesodermal cells and examined the role if TBBPA in disruption of mesoderm and mesoderm-derived blood cells. Our data shows that TBBPA exposures induced concentration-dependent decreases of Tbx16 protein for 0.75-8 hpf (p<0.0001) **(Figure 9A)** and 6-8 hpf (p<0.0001) **(Figure 9B)**. This suggests a disruption in mesodermal development and was consistent with our prior sequencing data on mRNA seq on 0.75 - 5 hpf of TBBPA exposure on zebrafish **(Table 1)**.

**Figure 9.**
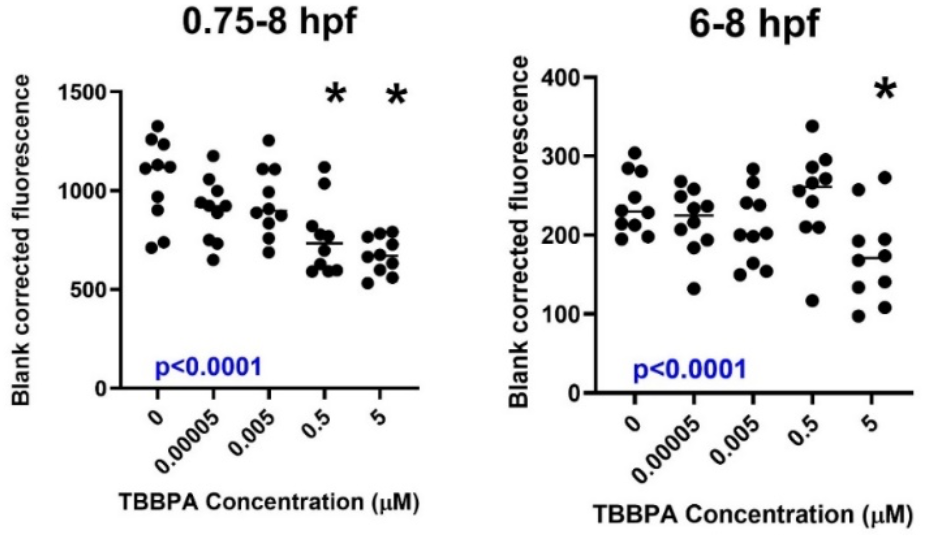
TBBPA decreases Tbx16 levels at (A) 0.75-8 hpf and (B) 6-8 hpf. Overall 1-way ANOVA p values are denoted by blue text; asterisk (*) denotes statistically different (p<0.05) from 0 µM based on Dunnett’s post hoc test following 1-way ANOVA.

We then quantified the impacts of TBBPA on blood cell production using o-dianisidine-a dye that stains hemoglobin. Our results show that TBBPA exposures induced concentration-dependent decreases in hemoglobin production area from 0.75 hpf to 3 dpf (p<0.0001) **(Figure 10)**, resulting in anemic embryos at concentrations down to 0.005 µM. Taken together, these suggest the potential for TBBPA to cause significant developmental issues in zebrafish, particularly in structures derived from the mesoderm where how TBBPA might disrupt key developmental processes, which is important for understanding their effects on embryonic development.

**Figure 10.**
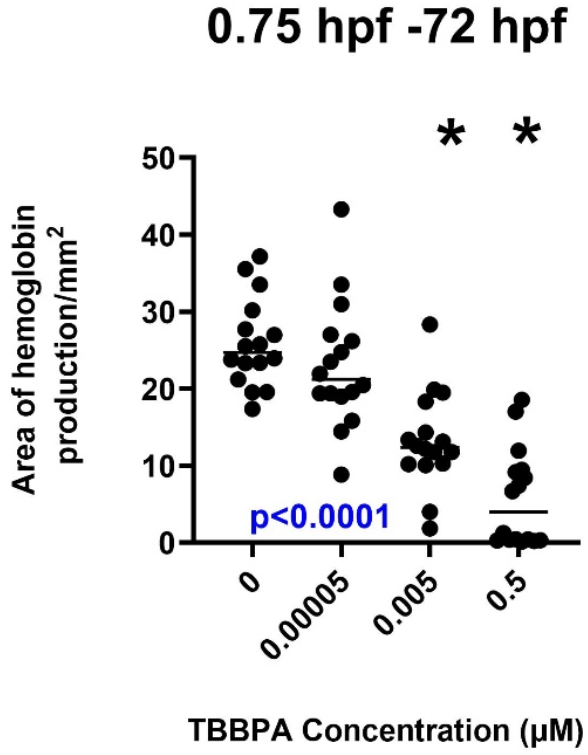
TBBPA decreases area of hemoglobin production at 0.75 hpf-72 hpf. Overall, 1-way ANOVA p values are denoted by blue text; asterisk (*) denotes statistically different (p<0.05) from 0 µM based on Dunnett’s post hoc test following 1-way ANOVA.

## Conclusion

Our data collectively indicates that exposure to TBBPA disrupts dorsoventral patterning by potentially targeting BMP signaling and EMT. Both these molecular phenomena are gatekeepers of downstream tissue specification, determination and differentiation. We further show that the integrity of neural crest cells and germ layers are disrupted on TBBPA exposures and that target cell types, such as cartilages and blood cells are impacted downstream. Collectively, we show that exposure to TBBPA has a systemic impact on early development. Importantly, many of these changes in molecular markers are seen at environmentally relevant concentrations, strengthening the importance of examining the effects of this ubiquitous flame retardants. Future studies are aimed at understanding the roles of specific gene products as mechanisms of these toxic effects.

## Acknowledgements

We acknowledge Mr. Scott Horman of Aquatic Animal Research Lab, Clemson University, for zebrafish husbandry.

## References

Alzualde, A., Behl, M., Sipes, N.S., Hsieh, J.-H., Alday, A., Tice, R.R., Paules, R.S., Muriana, A., Quevedo, C., 2018. Toxicity profiling of flame retardants in zebrafish embryos using a battery of assays for developmental toxicity, neurotoxicity, cardiotoxicity and hepatotoxicity toward human relevance. Neurotoxicol. Teratol. 70, 40–50. 10.1016/j.ntt.2018.10.002

Cariou, R., Antignac, J.-P., Zalko, D., Berrebi, A., Cravedi, J.-P., Maume, D., Marchand, P., Monteau, F., Riu, A., Andre, F., Bizec, B.L., 2008. Exposure assessment of French women and their newborns to tetrabromobisphenol-A: Occurrence measurements in maternal adipose tissue, serum, breast milk and cord serum. Chemosphere 73, 1036–1041. 10.1016/j.chemosphere.2008.07.084

Chen, J., Tanguay, R.L., Xiao, Y., Haggard, D.E., Ge, X., Jia, Y., Zheng, Y., Dong, Q., Huang, C., Lin, K., 2016. TBBPA exposure during a sensitive developmental window produces neurobehavioral changes in larval zebrafish. Environ. Pollut. 216, 53–63. 10.1016/j.envpol.2016.05.059

Dasgupta, S., Cheng, V., Vliet, S.M.F., Mitchell, C.A., Volz, D.C., 2018. Tris(1,3-dichloro-2-propyl) Phosphate Exposure During the Early-Blastula Stage Alters the Normal Trajectory of Zebrafish Embryogenesis. Environ. Sci. Technol. 52, 10820–10828. 10.1021/acs.est.8b03730

Dasgupta, S., Cheng, V., Volz, D.C., 2021. Utilizing Zebrafish Embryos to Reveal Disruptions in Dorsoventral Patterning. Curr. Protoc. 1, e179. 10.1002/cpz1.179

Dasgupta, S., LaDu, J.K., Garcia, G.R., Li, S., Tomono-Duval, K., Rericha, Y., Huang, L., Tanguay, R.L., 2023. A CRISPR-Cas9 mutation in sox9b long intergenic noncoding RNA (slincR) affects zebrafish development, behavior, and regeneration. Toxicol. Sci. 194, 153–166. 10.1093/toxsci/kfad050

Deng, Z.H., Li, Y.S., Gao, X., Lei, G.H., Huard, J., 2018. Bone morphogenetic proteins for articular cartilage regeneration. Osteoarthritis Cartilage 26, 1153–1161. 10.1016/j.joca.2018.03.007

Frey, P., Devisme, A., Schrempp, M., Andrieux, G., Boerries, M., Hecht, A., 2020. Canonical BMP Signaling Executes Epithelial-Mesenchymal Transition Downstream of SNAIL1. Cancers 12, 1019. 10.3390/cancers12041019

Gomez-Puerto, M.C., Iyengar, P.V., García De Vinuesa, A., Ten Dijke, P., Sanchez-Duffhues, G., 2019. Bone morphogenetic protein receptor signal transduction in human disease. J. Pathol. 247, 9–20. 10.1002/path.5170

Ho, K.-L., Yuen, K.-K., Yau, M.-S., Murphy, M.B., Wan, Y., Fong, B.M.-W., Tam, S., Giesy, J.P., Leung, K.S.-Y., Lam, M.H.-W., 2017. Glucuronide and sulfate conjugates of tetrabromobisphenol A (TBBPA): Chemical synthesis and correlation between their urinary levels and plasma TBBPA content in voluntary human donors. Environ. Int. 98, 46–53. 10.1016/j.envint.2016.09.018

Kalluri, R., Weinberg, R.A., 2009. The basics of epithelial-mesenchymal transition. J. Clin. Invest. 119, 1420–1428. 10.1172/JCI39104

Kim, Y.-S., Yi, B.-R., Kim, N.-H., Choi, K.-C., 2014. Role of the epithelial–mesenchymal transition and its effects on embryonic stem cells. Exp. Mol. Med. 46, e108–e108. 10.1038/emm.2014.44

Klymkowsky, M., Cortez Rossi, C., Artinger, K.B., 2010. Mechanisms driving neural crest induction and migration in the zebrafish and Xenopus laevis. Cell Adhes. Migr. 4, 595–608. 10.4161/cam.4.4.12962

Koehler, K.R., Malone, A.K., Hashino, E., 2014. Recapitulating Inner Ear Development with Pluripotent Stem Cells, in: Development of Auditory and Vestibular Systems. Elsevier, pp. 213– 247. 10.1016/B978-0-12-408088-1.00008-7

Langdon, Y.G., Mullins, M.C., 2011. Maternal and Zygotic Control of Zebrafish Dorsoventral Axial Patterning. Annu. Rev. Genet. 45, 357–377. 10.1146/annurev-genet-110410-132517

Li, S., Yang, R., Yin, N., Zhao, M., Zhang, S., Faiola, F., 2023. Developmental toxicity assessments for TBBPA and its commonly used analogs with a human embryonic stem cell liver differentiation model. Chemosphere 310, 136924. 10.1016/j.chemosphere.2022.136924

Li, Z., Jia, K., Chen, X., Guo, J., Zheng, Z., Chen, W., Peng, Y., Yang, Y., Lu, H., Yang, J., 2023. Exposure to Butylparaben Induces Craniofacial Bone Developmental Toxicity in Zebrafish (Danio rerio) Embryos. Ecotoxicol. Environ. Saf. 265, 115523. 10.1016/j.ecoenv.2023.115523

Liu, S., Deng, X., Zhou, X., Bai, L., 2021. Assessing the toxicity of three “inert” herbicide safeners toward Danio rerio: Effects on embryos development. Ecotoxicol. Environ. Saf. 207, 111576. 10.1016/j.ecoenv.2020.111576

Loh, C.-Y., Chai, J., Tang, T., Wong, W., Sethi, G., Shanmugam, M., Chong, P., Looi, C., 2019. The E-Cadherin and N-Cadherin Switch in Epithelial-to-Mesenchymal Transition: Signaling, Therapeutic Implications, and Challenges. Cells 8, 1118. 10.3390/cells8101118

Mahadeva, M., Niestępski, S., Kowacz, M., 2025. Modifying membrane potential synchronously controls the somite’s formation periodicity and growth. Dev. Biol. 517, 317–326. 10.1016/j.ydbio.2024.11.002

Miao, B., Yakubu, S., Zhu, Q., Issaka, E., Zhang, Y., Adams, M., 2023. A Review on Tetrabromobisphenol A: Human Biomonitoring, Toxicity, Detection and Treatment in the Environment. Molecules 28, 2505. 10.3390/molecules28062505

Morrow, Z.T., Maxwell, A.M., Hoshijima, K., Talbot, J.C., Grunwald, D.J., Amacher, S.L., 2017. tbx6l and tbx16 are redundantly required for posterior paraxial mesoderm formation during zebrafish embryogenesis. Dev. Dyn. 246, 759–769. 10.1002/dvdy.24547

Nifuji, A., Kellermann, O., Kuboki, Y., Wozney, J.M., Noda, M., 1997. Perturbation of BMP Signaling in Somitogenesis Resulted in Vertebral and Rib Malformations in the Axial Skeletal Formation. J. Bone Miner. Res. 12, 332–342. 10.1359/jbmr.1997.12.3.332

Ogata, S., Morokuma, J., Hayata, T., Kolle, G., Niehrs, C., Ueno, N., Cho, K.W.Y., 2007. TGF-β signaling-mediated morphogenesis: modulation of cell adhesion via cadherin endocytosis. Genes Dev. 21, 1817–1831. 10.1101/gad.1541807

Pang, S., Gao, Y., Wang, Y., Yao, X., Cao, M., Liang, Y., Song, M., Jiang, G., 2022. Tetrabromobisphenol A perturbs cell fate decisions via BMP signaling in the early embryonic development of zebrafish. J. Hazard. Mater. 430, 128512. 10.1016/j.jhazmat.2022.128512

Pomreinke, A.P., Soh, G.H., Rogers, K.W., Bergmann, J.K., Bläßle, A.J., Müller, P., 2017. Dynamics of BMP signaling and distribution during zebrafish dorsal-ventral patterning. eLife 6, e25861. 10.7554/eLife.25861

Ramel, M.-C., Hill, C.S., 2013. The ventral to dorsal BMP activity gradient in the early zebrafish embryo is determined by graded expression of BMP ligands. Dev. Biol. 378, 170–182. 10.1016/j.ydbio.2013.03.003

Serradimigni, R., Rojas, A., Leong, C., Pal, U., Bryan, M., Sharma, S., Dasgupta, S., 2024. Flame retardant tetrabromobisphenol A (TBBPA) disrupts histone acetylation during zebrafish maternal-to-zygotic transition. 10.1101/2024.03.31.587433

Shi, Z.-X., Wu, Y.-N., Li, J.-G., Zhao, Y.-F., Feng, J.-F., 2009. Dietary Exposure Assessment of Chinese Adults and Nursing Infants to Tetrabromobisphenol-A and Hexabromocyclododecanes: Occurrence Measurements in Foods and Human Milk. Environ. Sci. Technol. 43, 4314–4319. 10.1021/es8035626

Veldman, M.B., Lin, S., 2008. Zebrafish as a Developmental Model Organism for Pediatric Research. Pediatr. Res. 64, 470–476. 10.1203/PDR.0b013e318186e609

Wang, R.N., Green, J., Wang, Z., Deng, Y., Qiao, M., Peabody, M., Zhang, Q., Ye, J., Yan, Z., Denduluri, S., Idowu, O., Li, M., Shen, C., Hu, A., Haydon, R.C., Kang, R., Mok, J., Lee, M.J., Luu, H.L., Shi, L.L., 2014. Bone Morphogenetic Protein (BMP) signaling in development and human diseases. Genes Dis. 1, 87–105. 10.1016/j.gendis.2014.07.005

Zhao, M., Yin, N., Yang, R., Li, S., Zhang, S., Faiola, F., 2022. Environmentally relevant exposure to TBBPA and its analogues may not drastically affect human early cardiac development. Environ. Pollut. 306, 119467. 10.1016/j.envpol.2022.119467

Zhu, B., Zhao, G., Yang, L., Zhou, B., 2018. Tetrabromobisphenol A caused neurodevelopmental toxicity via disrupting thyroid hormones in zebrafish larvae. Chemosphere 197, 353–361. 10.1016/j.chemosphere.2018.01.080

